# Inhibitors of HIV-1 maturation modulate conformational dynamics and heterogeneity of the immature Gag lattice

**DOI:** 10.64898/2026.09.11.751012

**Authors:** Tathagata Nandi, Matthew Wong, Abdul A. Waheed, Eric O. Freed, Celia A. Schiffer, James B. Munro

## Abstract

HIV-1 maturation requires the viral protease to cleave the Gag polyprotein into its constituent domains. Subsequent conformational and morphological changes result in a mature, infectious virion. Prior studies support a hypothesis in which Gag conformational dynamics and the heterogeneity of the immature Gag lattice enable proteolytic and morphological maturation. We developed a single-molecule Förster resonance energy transfer imaging approach to probe, in real time, the conformational dynamics of individual Gag molecules within immature HIV-1 virions. Our results capture Gag’s conformational landscape and the spatial heterogeneity of the immature lattice. We evaluated inhibitors of maturation to identify immature lattice features that are modulated to disrupt maturation. Maturation inhibitors that act through diverse mechanisms, including lenacapavir and compounds that alter viral membrane composition, arrest Gag dynamics and reduce the spatial heterogeneity of the lattice. These observations support a model in which Gag dynamics and immature lattice organization regulate HIV-1 maturation.

## Introduction

HIV-1 is released from infected cells in an immature, non-infectious state. Virion maturation is required for downstream infection^1–4^. The major structural component of HIV-1 is the Gag polyprotein, which consists of the matrix (MA), capsid (CA), nucleocapsid (NC), and p6 domains, as well as spacer peptides SP1 and SP2^1,3,5^. Thousands of Gag monomers assemble to form a lattice within the immature virion^6–8^. The immature lattice is composed of 95% Gag and 5% GagPol, which contains the viral protease PR^9^. For PR to function, GagPol must dimerize, which may require diffusion within the immature lattice^9,10^. Maturation involves a cascade of temporally regulated proteolysis events, in which the Gag monomers are cleaved into their constituent domains by PR^10–12^. These cleavage events occur at distinct rates, with CA-SP1 cleavage being the slowest and rate-limiting for maturation^5,6^. Therefore, PR must access different cleavage sites across the immature Gag lattice, implicating the need for lattice dynamics^8,9^. Inhibiting cleavage or interfering with the order of cleavage disrupts maturation, preventing downstream infection, which is a proven therapeutic strategy^13,14^. Following proteolysis, the Gag domains undergo conformational and organizational changes that result in the formation of the mature, infectious virion. Targeting Gag to inhibit maturation is an emerging strategy for therapeutic development.

The immature Gag lattice assembles as an incomplete sphere with a large gap at the budding site and irregularly shaped defects in the hexagonal array^8,15–18^. The MA domain forms hexamers of trimers^4,19^, whereas the CA domain forms hexamers of monomers^1,6,8^. The different arrays in the MA and CA layers indicate structural flexibility of Gag within the lattice. Only one- to two-thirds of the lattice contains well-ordered hexagonal Gag^9,17,18^. Gag monomers at the lattice peripheries can form partial hexamers with fewer stabilizing intermolecular contacts, potentially enabling greater flexibility^8,9^. These structural features of the immature lattice implicate Gag conformational heterogeneity and dynamics that have not been observed experimentally. We hypothesize that lattice heterogeneity and Gag flexibility are important for PR dimerization and activation, and that they further enable PR to access cleavage sites and facilitate the morphological changes needed for maturation^20–22^. To test this hypothesis, we developed an experimental method centered on single-molecule Förster resonance energy transfer (smFRET) imaging, which enables visualization of the conformational dynamics of individual Gag molecules within the immature virion and captures their spatial heterogeneity.

Maturation inhibition is a therapeutic strategy originally implemented by targeting PR^13^. Targeting Gag later emerged as a viable alternative^2,14^. Bevirimat (BVM) targets the CA-SP1 cleavage site by preventing the localized dynamics needed to expose it^23–25^. The T8I mutation in SP1 phenocopies the effect of BVM and results in further strengthening of the CA-SP1 six-helix bundle in the immature lattice^24,26–28^. Lenacapavir (LEN) is an FDA-approved, first-in-class inhibitor that binds both mature and immature CA and affects multiple steps in the HIV-1 replication cycle^29–32^. LEN promotes assembly of immature virions with aberrant topology, suggesting modulation of Gag conformation and intermolecular contacts, which reduces mature, infectious virions^33^. More recently, targeting the cellular enzyme neutral sphingomyelinase 2 (nSMase2), which disrupts the balance of sphingomyelin and ceramide in the viral membrane, emerged as a novel strategy for inhibiting HIV-1 maturation^34,35^. Targeting nSMase2 in virus-producing cells reduced Gag proteolytic processing, despite no effect on Gag or GagPol incorporation, suggesting either inhibited PR activation or reduced access to cleavage sites^34,35^. In all cases, Gag conformational dynamics within the immature lattice are implicated in the mechanism of inhibitor action.

Here, we have used smFRET imaging to reveal the conformational dynamics of individual Gag proteins within the immature virion. With this approach, we tested the hypothesis that Gag conformational dynamics within the immature lattice guide proteolysis and maturation^17^. Our data indicate a dynamic equilibrium among multiple conformational states. Classification of the smFRET trajectories based on dynamic behavior revealed Gag in well-ordered regions of the immature lattice, at lattice edges, and as monomers or small assemblies free from significant structural constraints. We find that maturation inhibitors modulate Gag conformational dynamics and lattice heterogeneity in manners that inform on their mechanisms of action. Notably, our data show that LEN arrests Gag conformational dynamics and reduces spatial heterogeneity of the immature lattice. Inhibition of nSMase2 during virion production also reduced Gag conformational dynamics and heterogeneity, suggesting that two different pathways can disrupt maturation by affecting the immature lattice. These results indicate that Gag conformational dynamics and immature lattice spatial heterogeneity regulate proteolysis and subsequent maturation.

## Results

### Site-specific labeling of Gag inside HIV-1 virions

Prior studies demonstrated that CA-SP1 cleavage is incomplete when MA-CA cleavage is restricted^5^. Furthermore, formation of the mature conical capsid requires liberating CA from MA^3,5^. These observations underscore the importance of studying the dynamics across the MA– CA interface within intact immature virions. However, the flexible linker between the MA and CA layers has limited structural description^19^. Therefore, to develop an smFRET imaging assay, we site-specifically labeled the MA and CA domains to probe their relative dynamics within intact immature virions. To attach fluorophores to Gag, we used a genetic code expansion approach^36,37^. We introduced an amber stop codon (TAG) at position 92 in the loop between helix 5 and 6 of MA (MA92*), which avoided trimer interfaces, the highly basic region, and envelope interaction sites^38,39^. We introduced a second amber stop codon at position 95 within the cyclophilin-A (CypA) loop of the NTD of CA (CA95*). We made these modifications in a *pol*-deleted pNL4-3 construct. Amber codon readthrough incorporated a non-natural amino acid containing a trans-cyclooctene moiety (Fig. S1a). We predicted that this modification would not alter immature virus production, as CypA binding is restricted to residues 85-93 and is not required for virus assembly^40^. Virions formed with Gag^MA92*/CA95^* had no loss of proteolytic maturation as compared to wild-type Gag (Fig. S1b). Negative-stain electron microscopy and analytical density gradient ultracentrifugation showed that immature HIV-1^MA92*-CA95^* virions were of comparable size and morphology to wild-type virions (Fig. S3). Membrane-permeable tetrazene-conjugated donor and acceptor fluorophores (JF549 and JF646) were then site-specifically attached to MA92* and CA95* inside immature virions using copper-free click chemistry. TIRF microscopy and steady-state fluorescence demonstrated site-specific fluorescent labeling (Fig. S2). Based on molecular dynamics (MD) simulation of a MA-CA 18-mer model generated using AlphaFold and previously reported structures (see Materials and Methods), we estimated the distance between fluorophores attached at MA92* and CA95* to be *ca.* 60 Å (Fig. 1b). MD simulation showed a broad distribution of distances when applied to the MA-CA monomer. These models suggest that the MA and CA domains undergo significant relative movement within the Gag monomer but prefer an extended conformation within the hexameric immature lattice (Fig. 1b).

**Fig. 1.**
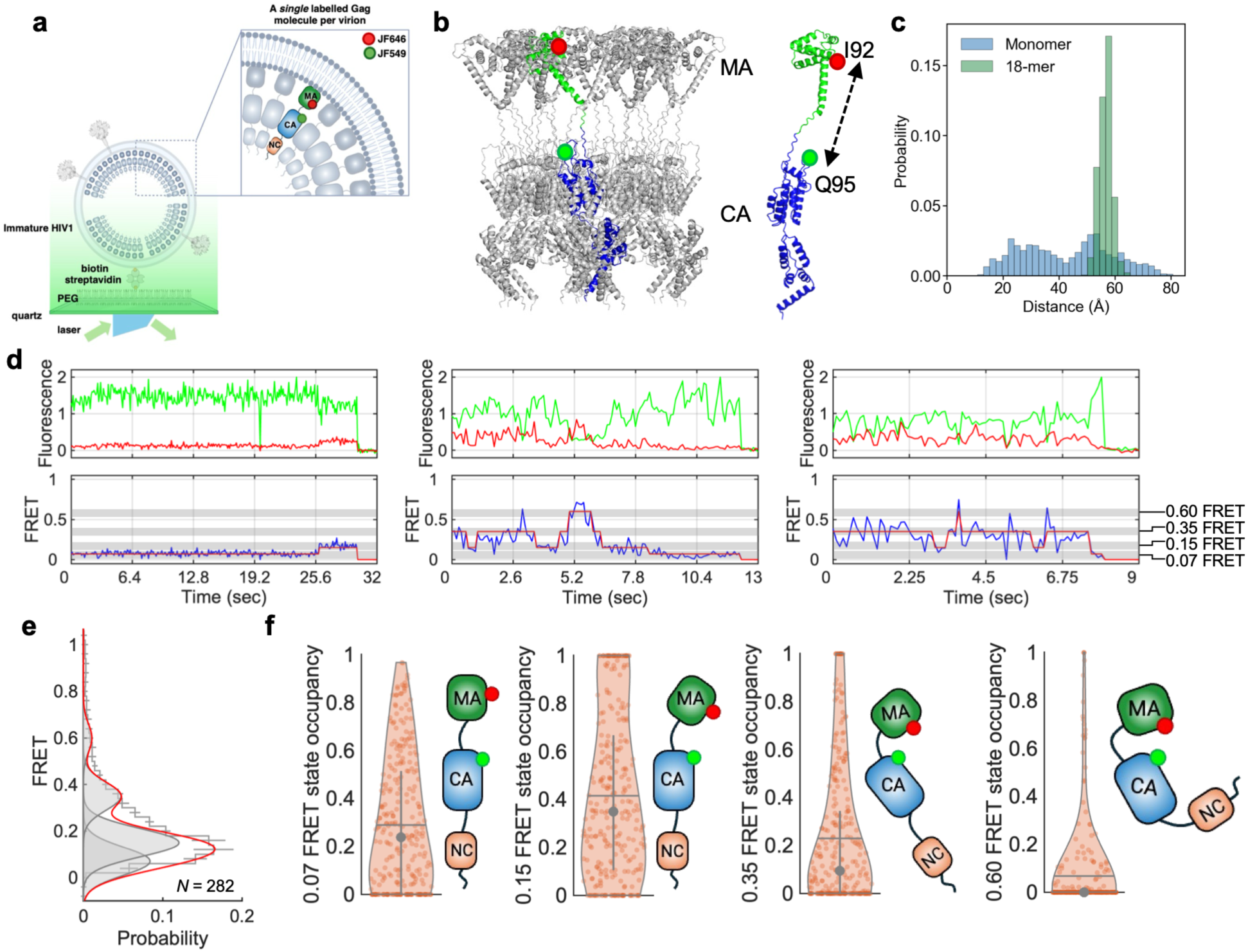
Gag conformational dynamics within the immature HIV-1 virion. (**a**) smFRET imaging using prism-based TIRF microscopy with an immobilized immature virion containing one Gag protein fluorescently labeled with JF549 (green) and JF646 (red). (**b**) Atomic models of an 18-mer assembly of MA-CA with one fluorescently labeled Gag monomer highlighted (MA, green; CA, blue), and a MA-CA monomer. Fluorophore attachment sites are indicated. (**c**) Distributions of inter-fluorophore distances determined by MD simulations of the MA-CA monomer and 18-mer. (**d**) Three representative smFRET trajectories showing donor (green) and acceptor (red) fluorescence and corresponding FRET trajectories (blue). The four non-zero FRET states identified through HMM analysis are indicated with grey shading. The idealizations resulting from HMM analysis are overlaid on the FRET trajectories (red). (**e**) FRET histogram compiled from 282 smFRET trajectories, with overlaid Gaussian distributions representing the 4 non-zero FRET states identified through HMM. Histogram bins reflect the mean of three pools of trajectories, with error bars indicating the standard error. (**f**) Violin plots representing the distributions of occupancies in each of the four non-zero FRET states obtained from 282 molecules. The grey horizontal lines indicate the mean occupancies; the grey circles indicate the median; and the vertical lines represent the 25-75% quartiles.

### Gag is conformationally dynamic inside the virion

smFRET imaging required virions in which a single Gag monomer is labeled with one pair of donor and acceptor dyes within the immature lattice (Fig. 1a). We therefore produced immature virions with a large excess of wild-type HIV-1 plasmid over HIV-1^MA92*-CA95^* plasmid and performed site-specific labeling as described above. We empirically determined the plasmid ratio to obtain fluorescence traces from single Gag proteins within virions immobilized on a quartz microscope slide and imaged with TIRF microscopy (Fig. 1c). The FRET efficiency reflects Gag conformation in terms of the relative positions of MA and CA. The trajectories indicated multiple FRET states, suggesting that Gag exists in distinct conformations within the immature virion, which are in dynamic equilibrium. Application of hidden Markov modeling (HMM) to the FRET trajectories identified 4 non-zero FRET efficiencies (0.07 ± 0.03, 0.15 ± 0.05, 0.35 ± 0.09, and 0.6 ± 0.1), and a 0-FRET state corresponding to photobleached fluorophores (Fig. 1d). Histograms of FRET efficiencies show that the 0.07- and the 0.15-FRET states are thermodynamically most stable (Fig. 1d, e). Occupancy violin plots represent the fraction of time each smFRET trajectory spends in each of the 4 non-zero FRET states (Fig. 1f). The FRET traces displayed 29 ± 2% and 42 ± 2% mean occupancy in the 0.07- and 0.15-FRET states, respectively (Fig. 1d, e). The low FRET efficiencies of these states suggest extended Gag conformations within the lattice. The transition density plot (TDP) indicates the frequency of transitions between the FRET states (Fig. 2c). The most frequent transitions are between the 0.07- and 0.15-FRET states, which suggests a dynamic equilibrium among structurally similar extended Gag conformations. Analysis of the kinetics of transitions between the FRET states showed that the 0.07- and 0.15-FRET states are long-lived relative to the higher-FRET states, suggesting Gag conformations with MA and CA stabilized by intermolecular contacts within the lattice (Fig. 2b, Fig. S5). The 0.35- and 0.6-FRET states may reflect more compact Gag conformations in which MA and CA become more proximal. The 0.35-FRET state is accessible from both lower FRET states. But 0.35 FRET is adopted more frequently by way of 0.15 FRET, suggesting that 0.15 FRET represents an intermediate Gag conformation between 0.07 and 0.35 FRET (Fig. 2c). We rarely observed transitions into or out of the 0.6-FRET state, suggesting a Gag conformation that is accessible in a distinct context. Taken together, these data indicate a dynamic description of Gag conformation within the immature HIV- 1 virion, in which extended and compact conformations are accessible.

**Fig. 2.**
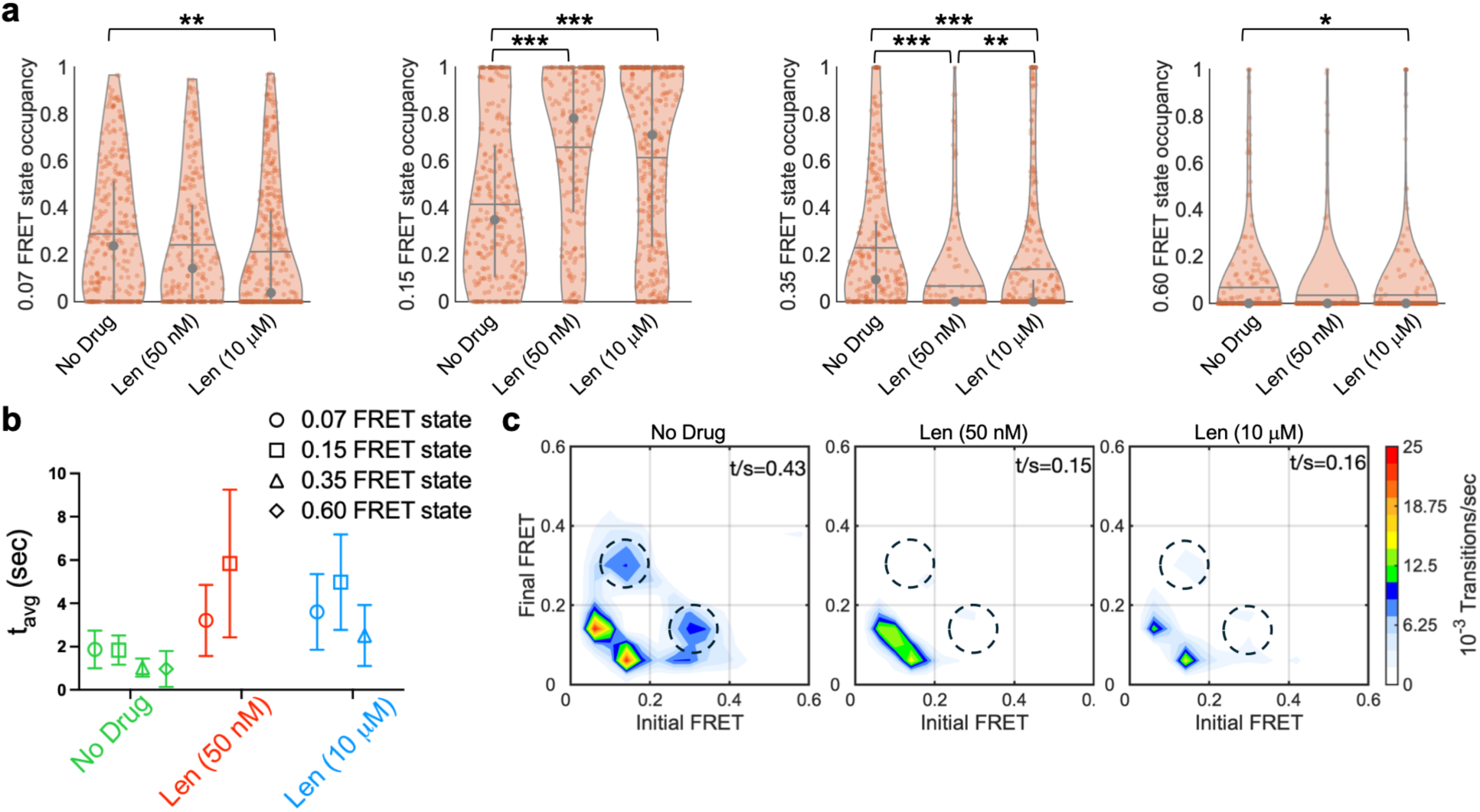
LEN decreases the conformational dynamics of Gag in the immature HIV-1 virion. (**a**) Occupancy violin plots acquired in the absence of inhibitor (282 molecules); in the presence of 50 nM LEN during virion production (231 molecules); and in the presence of 10 μM LEN added to purified virions (395 molecules). Violin plots are displayed as in Fig. 1. Asterisks indicate the statistical significance: ***, *p* < 0.001; **, *p* < 0.01; *, *p* < 0.05; the absence of an asterisk indicates no statistical significance. (**b**) Average dwell time (*t*_avg_) of each FRET state for each of the three conditions (determined in Fig. S5). Error bars indicate the 95% confidence intervals. The absence of a *t*_avg_ indicates it was not determined due to insufficient observation of dwells. (**c**) TDPs indicating the reduction of Gag conformational dynamics in the presence of LEN. *t*/*s* indicates the total transitions observed per second. Dashed circles indicate transitions into and out of the 0.35- FRET state, which are abrogated in presence of LEN.

### Lenacapavir reduces Gag conformational dynamics

We next tested the hypothesis that Gag conformational signatures within the immature virion, which are reflected in our smFRET data, are relevant to maturation. LEN binds the NTD of CA at the hexameric interface and rigidifies the immature lattice, disrupting maturation^31,33^. We introduced LEN to virions in two ways before smFRET imaging. First, we produced immature virions in the presence of 50 nM LEN. Second, we produced immature virions without LEN, followed by incubation with 10 μM Len for 1 hr at room temperature. In both cases, we observed the same four FRET states as seen in the absence of LEN but with altered occupancies. LEN shifted the conformational equilibrium such that 0.15 FRET had the highest occupancy (Fig. 2a, Fig. S4) and longest lifetime (Fig. S5). The 0.35- and 0.6-FRET state occupancies were reduced significantly (Fig. 2a, Fig. S4). The TDP indicated that both LEN treatments reduced conformational transitions, specifically abrogating transitions to and from the 0.35-FRET state (Fig. 2c). Consistently, the average lifetimes of 0.07- and 0.15-FRET states were increased (Fig. 2b). Overall, the effect of LEN was greater when added during virion assembly. This suggests that adding LEN to virus-producing cells affects immature lattice formation, which in turn shapes the Gag conformational landscape. We interpret these observations as LEN stabilizing an extended Gag conformation, reducing compact conformations, and limiting overall dynamics within the lattice. This interpretation is consistent with structural and cellular imaging data indicating a flattened and rigidified immature Gag lattice in the presence of LEN^33^. Hence, our smFRET data demonstrate the structural and dynamic properties of Gag within the immature virion that are relevant to downstream maturation.

### Allosteric control of MA-CA motions

We next asked whether our smFRET data report on the stability of the CA-SP1 helical bundle, which is critical for proteolysis and maturation. BVM, which inhibits proteolysis of the CA-SP1 cleavage site, only showed a slight increase in the 0.07-FRET state (Fig. S6). The T8I mutation in SP1, which stabilizes the CA-SP1 helical bundle^24,26,28^, increased the occupancy in the 0.07-FRET state to a greater extent than BVM. T8I further reduced occupancy of the 0.35- and 0.60-FRET states (Fig. S6). Together, these results suggest that the 0.07-FRET state, which is consistent with linear Gag conformation, reflects a stable CA-SP1 helical bundle. The relatively modest effect of BVM on FRET-state occupancies likely reflects the fluorophore locations, which are distal to the BVM-binding site at the CA-SP1 junction. Both BVM and T8I slightly increased the frequency of transitions between the FRET states as observed in their TDPs (Fig. S6b). This suggests that although BVM and T8I allosterically mediate Gag conformation, they do not arrest the relative motions between MA and CA.

### Viral membrane composition controls Gag conformation

Next, we investigated the effect of nSMase2 inhibitors on Gag conformation to understand better how disruption of viral membrane composition inhibits maturation. We produced virions in the presence of 10 μM PDDC, an nSMase2 inhibitor, and performed smFRET experiments as above. PDDC increased the 0.15 FRET occupancy to 48 ± 2%, compared with 42 ± 2% in the absence of the inhibitor. PDDC also decreased the 0.35 FRET occupancy to 14 ± 1% (Fig. 3a), compared with 23 ± 1% in the absence of the inhibitor. The lifetimes of 0.07 and 0.15 FRET states increased in the presence of PDDC, which reduced the frequency of transitions to 0.35 FRET (Fig. 3b-c, Fig. S5). Similar but milder effects were observed for another nSMase2 inhibitor, DPTIP (Fig. S8). The reduction in transition frequencies implies that the depletion of ceramides or enrichment of sphingomyelin in the viral membrane resulting from nSMase2 inhibition leads to a more ordered immature lattice with reduced dynamics, which disrupt proteolysis (Fig. S7). 10 μM LEN added to purified immature virions produced in the presence of PDDC had an additive effect on Gag dynamics, resulting in the greatest stabilization of 0.15 FRET, observed both as increased occupancy (69 ± 1%) and lifetime (Fig. 3). The additive effect indicates that LEN and PDDC function by independent mechanisms to modulate the immature Gag lattice properties and disrupt maturation. These data underscore the correlation between stabilization of linear Gag conformations, reduced overall lattice dynamics, and maturation inhibition. Thus, our results support a model in which HIV-1 virion maturation requires flexibility in Gag conformation within the immature lattice.

**Fig. 3.**
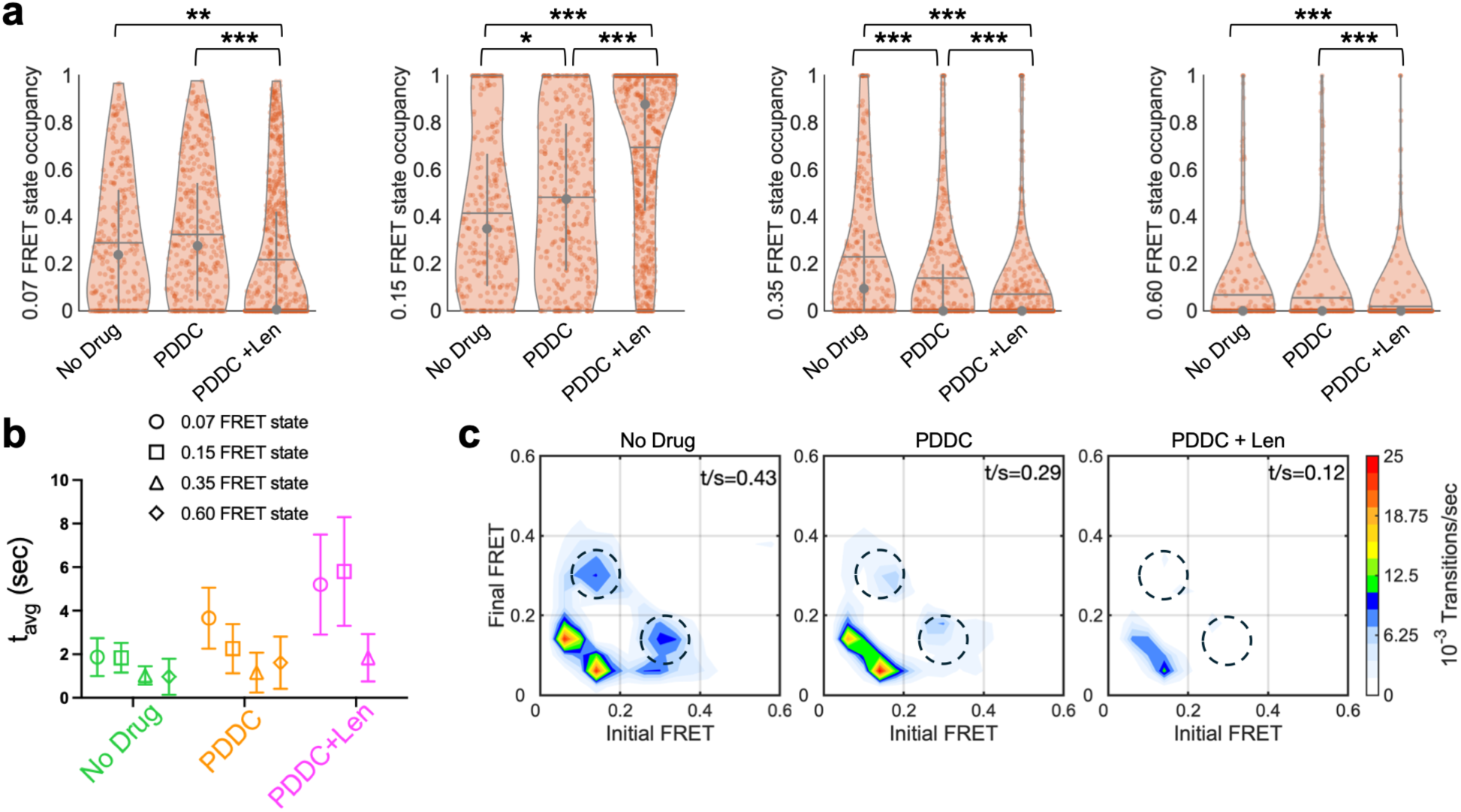
Disruption of viral membrane content reduces HIV-1 Gag conformational dynamics. (**a**) Occupancy violin plots displayed as in Fig. 2. The data were acquired from virions produced in the absence (282 molecules) or presence of PDDC (369 molecules), and in the absence and presence of 10 μM LEN (659 molecules) added to the purified virions. (**b**) Average dwell time (*t*_avg_) of each FRET state for each of the three conditions, displayed as in Fig. 2 (determined in Fig. S5). (**c**) TDPs indicating the reduction of Gag conformational dynamics in the presence of PDDC, and in the added presence of LEN. *t*/*s* indicates the total transitions observed per second. Dashed circles indicate transitions into and out of the 0.35-FRET state, which are abrogated by PDDC and LEN.

### Classification of Gag dynamics indicates distinct environments within the virion

Visual inspection of FRET trajectories indicated three subpopulations. We asked whether these subpopulations could be correlated with distinct lattice environments within the immature virion. We quantified these subpopulations using *k*-means clustering of the FRET trajectories based on combined occupancies of the 0.07- and 0.15-FRET states, which reflect extended Gag conformations, and the number of FRET transitions (Fig. S9). In the absence of inhibitors, Cluster 1 was the largest subpopulation, comprising 40% of the total. Cluster 1 contained trajectories with the highest combined occupancy in the low-FRET extended conformations and showed transitions only between these states (Fig. 4b, Fig. S10). Trajectories in Cluster 2 accessed the 0.35-FRET state, which reflects a more compact conformation, but still favored the low-FRET extended states (Fig. 4b). Trajectories in Cluster 3 showed the highest occupancy in the 0.35-FRET state, with frequent transitions to the low-FRET extended states and the 0.6-FRET state, which reflects further Gag compaction (Fig. 4b, Fig. S10). This suggests a progressive increase in conformational freedom from Cluster 1 to Cluster 2 to Cluster 3, enabling Gag to access both compact and extended conformations. Based on these observations, Cluster 1 likely represents the interior homogeneous regions of the immature Gag lattice, where complete Gag hexamers predominate, and Gag-Gag interactions stabilize the extended conformations. Consistent with this interpretation, structural data indicate that approximately 40% of the viral surface is covered with well-ordered hexagonal Gag, which agrees with the size of the Cluster 1 subpopulation determined here^9,17,18^. Cluster 2 may represent lattice edges where Gag can access compact conformations because fewer structural constraints apply within the partial hexamers. The hexameric symmetry likely still exists because the extended conformations are still favored over the compact conformations in Cluster 2. Finally, Cluster 3 may represent Gag molecules free from the hexameric lattice, allowing Gag greater access to compact conformations and enabling MA and CA to come into proximity. Isolated Gag monomers in solution favor compact conformations more than the extended conformation in the absence of assembly co-factors^41^.

**Fig. 4.**
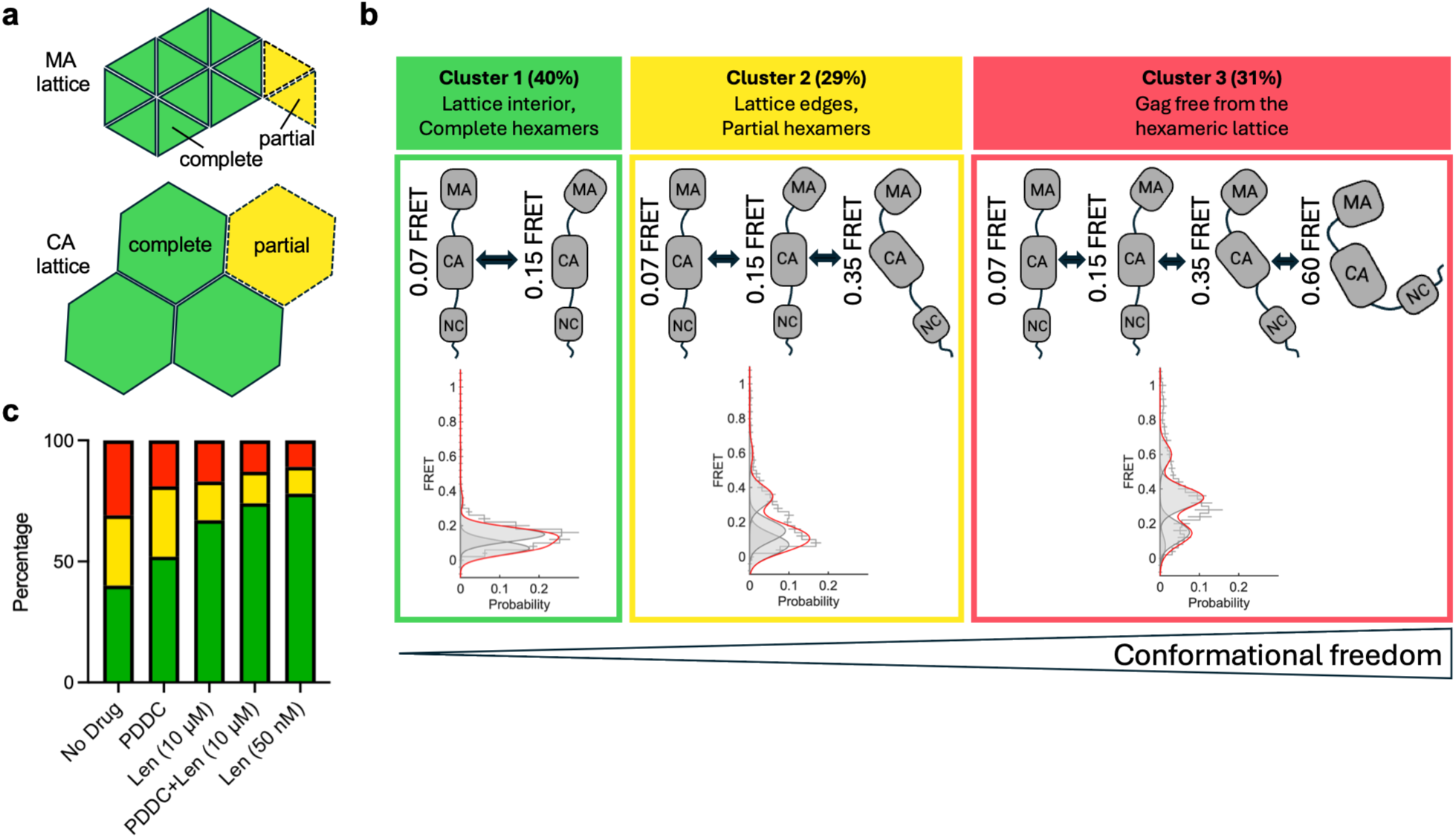
Clustering of smFRET trajectories indicates spatial heterogeneity within the immature HIV-1 virion. (**a**) Patch of immature Gag lattice showing hexamers of MA trimers and hexamers of CA monomers. Complete MA trimers and CA hexamers in the lattice interiors (Cluster 1) are indicated by green triangles and hexagons with solid borders, respectively. Incomplete or partial MA trimers and CA hexamers at the lattice edges (Cluster 2) are indicated by yellow triangles and hexagons with dashed borders. (**b**) Cartoons of available Gag conformations within each Cluster, with conformational freedom increasing from Cluster 1 to 2 to 3. Only FRET states with at least 5% occupancy for each cluster are shown. Also shown are FRET histograms for each Cluster obtained in the absence of inhibitor. (**c**) The Cluster distribution shown for different experimental conditions. Inhibitors reduce the spatial heterogeneity, increasing Cluster 1 and decreasing Clusters 2 and 3. Clusters are colored as in b.

We applied the same clustering algorithm to smFRET trajectories acquired in the presence of inhibitors. This analysis showed that conditions that inhibit maturation increased the subpopulation in Cluster 1 to 78% when virions were formed with 50 nM LEN and 67% when 10 μM LEN was added to purified virions, compared with 40% in the absence of inhibitors (Fig. 4c). Inhibition of nSMase2 by PDDC increased Cluster 1 to 52% (Fig. 4c). The combination of 10 μM LEN and PDDC increased Cluster 1 to 74% (Fig. 4c). These results are consistent with diverse maturation inhibitors stabilizing the hexameric lattice and reducing heterogeneities, which leads to loss of Gag conformational dynamics. In contrast, BVM and T8I did not show significant changes in cluster distribution, suggesting that their local CA-SP1 stabilizing effect is insufficient to impose global order in the immature Gag lattice with respect to MA-CA distances. This is consistent with structural data showing no significant change in Gag lattice structure in virus-like particles made in the presence of BVM or T8I^31^. Interestingly, BVM specifically inhibits the CA- SP1 cleavage during maturation, and its resistance mutations lie in the CA-SP1 junction^23,24,27,42,43^.

## Discussion

Our results demonstrate that HIV-1 Gag can access at least four different conformations with respect to MA-CA orientation, which are in a dynamic equilibrium within the immature lattice. Gag diffusion inside the virion, including attachment and detachment of Gag monomers within the immature lattice, may also influence our observations^44^. We identified Gag subpopulations based on their dynamic signatures, which correlate with distinct environments within the immature virion. The effects of diverse inhibitors support the relevance of these dynamics to maturation. Whether through direct interaction with Gag or disrupting viral membrane content, these inhibitors consistently reduce Gag conformational dynamics and lattice heterogeneity, as reflected in an increase in the Cluster 1 subpopulation. These observations support our hypothesis that Gag conformational dynamics and lattice heterogeneity are critical for virion maturation.

Multiple explanations exist for how Gag dynamics and lattice heterogeneity facilitate maturation. First, dynamics and heterogeneity may permit the viral PR to access the cleavage sites. For example, incomplete CA hexamers at lattice boundaries may destabilize the CA-SP1 helical bundle, enhancing transient exposure of the cleavage site^8,9^. Reducing such lattice heterogeneity would limit Gag flexibility at this functionally critical location, inhibiting proteolysis and maturation. Second, Gag flexibility and heterogeneity may enable disassembly of the immature lattice organization, which is necessary for reassembly of the mature capsid. Thermodynamic stabilization of the immature lattice may trap CA monomers following proteolysis in the immature geometry, disallowing the conformational changes needed for mature hexamer and pentamer formation. Finally, flexibility in the immature lattice may enable GagPol diffusion within the virion, which could facilitate dimerization and PR activation. Further investigations are needed to establish better the mechanistic link between viral membrane composition, Gag lattice heterogeneity and dynamics, and maturation.

Our results offer new insights into how Gag proteolysis, conformational changes in Gag, and morphological changes of the virion coalesce during HIV-1 maturation. Aberrant virion morphology has been reported when producer cells are treated with either LEN or PDDC. Immature virions formed in the presence of LEN exhibited a “mature-like” CA layer with flat facets that lacked the curvature of the native immature lattice^33^. While LEN may not affect proteolysis of immature Gag, it resulted in aberrant capsid morphologies that lacked the CA pentamers needed to form the mature conical capsid^33,45^. In light of these prior results, our data show that the lack of lattice curvature induced by LEN is commensurate with a loss of Gag conformational dynamics and reduced lattice heterogeneity. Furthermore, Gag flexibility is essential for CA monomers to reassemble into pentamers needed to form the conical mature capsid. Similarly, PDDC treatment disrupted the balance of sphingomyelin and ceramides in the viral membrane, resulting in immature virions with large gaps in the Gag lattice and membranous blebs protruding from the viral surface^35^. This may indicate consolidation of small discontinuities in the immature lattice into the observed large gaps, consistent with our observation of reduced dynamics and lattice heterogeneity, and the increase in Gag molecules found in Cluster 1. Our results show that both LEN and PDDC arrest immature Gag lattice dynamics and reduce spatial heterogeneity. However, PDDC treatment also disrupts Gag proteolysis (Fig. S7)^35^. This suggests that nSMase2 inhibitor treatment alters viral membrane composition in a way that arrests PR activation, leading to disruption of Gag proteolysis. In contrast, LEN directly interacts with Gag molecules and perturbs similar immature Gag lattice parameters, disrupting the structural and morphological aspects of maturation without affecting proteolysis^45^. This further supports the idea that Gag-membrane interactions mediate Gag mobility within the lattice, aiding maturation either by enabling GagPol dimerization, facilitating PR access to cleavage sites, or both. Taken together, these observations link viral membrane composition, virion morphology, Gag conformational dynamics, and lattice heterogeneity, informing the mechanism of maturation.

## Materials and methods

### Plasmid constructs

We have used pNL4-3 ΔRTΔPR (partially deleted reverse transcriptase (RT) and PR) as for production of non-infectious immature HIV-1 virions. For amber suppression, Gag was mutated at I92 and Q227 (95 in CA numbering) to TAG amber codons. The engineered construct, pNL4-3 ΔRTΔPR(I92TAG/Q227TAG) was obtained from Genscript and was used to produce HIV-1^MA92*-CA95^*. The T8I mutation was also introduced by Genscript into the same plasmid. For maturation assays, a plasmid expressing full-length HIV-1 Gag and GagPol under a CMV promoter (pGagPol) was used to produce proteolytically matured but non-infectious virions. Identical amber mutations were introduced into this plasmid.

### Mammalian cell culture

HEK293T cells were maintained at 37°C in DMEM (GIBCO) media supplemented with 100 U/ml penicillin/streptomycin (GIBCO), 2mM Glutamine (GIBCO), and 10% cosmic calf serum (CCS). Cells were passaged at 70% confluency and used for transfection and virion production.

### Modification of HIV-1 Gag to enable fluorescent labeling

The pNL4-3 ΔRTΔPR(I92TAG/Q227TAG) plasmid was tested for amber read through efficiency. HEK293T cells were seeded in six-well plates at 70% confluency and transfected with 4 μg pNL4-3 ΔRTΔPR or pNL4-3 ΔRTΔPR(I92TAG/Q227TAG). Polyethylenimine (PEI; Polyscience, Niles, IL) was used as the transfection reagent. For the amber-containing construct, the cells were also transfected with 0.4 μg of a plasmid encoding the orthogonal tRNA synthetase/tRNA (Mm-PylRS-AF/Pyl-tRNA^CUA^), and 0.4 μg of plasmid encoding eukaryotic translation termination factor 1 (eRF1) in presence of the non-natural amino acid trans-Cyclooct- 2-en – L - Lysine (TCO*A) (Sirius Fine Chemicals, Bremen, Germany). The orthogonal tRNA synthetase/tRNA and eRF1 ensured amber read through at I92 and Q227 and incorporation of TCO*A. The amount of full-length HIV-1 Gag (p55) obtained indicated the efficiency of amber read through. Virus was grown for 24 hrs at 37°C post-transfection. 1.5 ml of viral supernatant was collected and centrifuged over a 0.1-ml 10% sucrose cushion for 2 hours at 14000 rpm to concentrate the virions. The resulting pellets were resuspended in Laemmli sample buffer containing 2.5% b-mercaptoethanol and heated at 95°C for 10 mins before evaluation on mini-PROTEAN-TGX-4-20% precast gels (Biorad, Hercules, CA). The gels were used for Western blotting using a Trans-Blot Turbo (Biorad). Full-length HIV-1 Gag was probed with a murine p24 primary antibody (ThermoFisher Scientific, Waltham, MA), followed by an anti-mouse IgG-HRP secondary antibody, and developed with super signal west-pico plus chemiluminescent substrate (ThermoFisher Scientific). Fig. S1a shows the resulting Western blot, which indicated a 55 kDa band for full-length Gag in the presence of Mm-PylRS-AF/Pyl-tRNA^CUA^ and eRF1. This verifies positive amber read through with incorporation of TCO*A.

### Proteolytic maturation assay

HEK293T cells grown in six-well plates were transfected with 4 μg pGagPol, or 4 μg pGagPol(I92TAG/Q227TAG) with 0.4 μg Mm-PylRS-AF/Pyl-tRNA^CUA^ and eRF1 plasmids in the presence of TCO*A. PEI was used as the transfection reagent. Viruses were allowed to grow for 48 hrs post-transfection. Virions were collected and concentrated, and probed for p24 by Western blot, as described above. The presence of p24 indicated successful cleavage of full-length Gag. This analysis also shows that the amber mutations do not affect proteolysis by PR.

### Site-specific labelling of Gag within immature HIV-1 virions

Site-specific fluorophore attachment to Gag within the virion was performed using strain promoted inverse electron demand Diels Alder chemistry (SPIEDAC)-based click chemistry between TCO*A and tetrazene-conjugated JF549 (donor) and JF646 (acceptor) (Tocris Biosciences, Bristol, UK). Following transfection, as described above, virions were harvested from three 10-cm plates. Supernatants were filtered using 0.45 μm syringe filters and centrifuged over a 5 ml 10% sucrose cushion at 25000 rpm for 2 hrs at 4°C. The resulting viral pellet was resuspended in 600 μL PBS. The virus suspensions were incubated in the dark with 500 nM JF549- Tet and JF646-Tet for 1 hr at room temperature, followed by 30 mins at 37°C. The reaction mixtures were then treated with 0.2 mg DSPE-PEG-biotin resuspended in 40 μL PBS for an additional 30 mins at room temperature with gentle mixing. To remove unbound fluorophore, the mixtures were dialyzed into fresh PBS using 10 kDa Slide-A-Lyzer™ MINI Dialysis devices (ThermoFisher Scientific) for 2 hrs at room temperature, followed by a second round of dialysis overnight at 4°C. The dialyzed virions were then purified on a 12-mL 6-30 % opti-prep density gradient medium (Millipore Sigma, Burlington, MA) through ultracentrifugation at 35000 rpm for 1 hr at 4°C. Peak fractions were identified by p24 Western blot, as described above, pooled, and stored in aliquots at -80°C. The relative concentration of HIV-1^MA92*/CA95^* and wild-type HIV-1 was estimated from p55 intensities on Western blots using a dilution series of wild-type HIV-1 (Fig. S2a). We estimated wild-type HIV-1 to be *ca.* 800 times more concentrated than HIV- 1^MA92*/CA95^* due to low read through efficiency of the amber codons in pNL4-3 ΔRTΔPR (I92TAG/Q227TAG). Following equalization of concentration, we measured the bulk fluorescence intensity of fluorescently labelled HIV-1^MA92*/CA95^* (Fig. S2b). In contrast, wild-type mock-labeled HIV-1 showed only background fluorescence. The virions were also immobilized on streptavidin-coated quartz microscope slides and imaged with TIRF microscope (Fig. S2c). Consistent with the bulk measurements, only HIV-1^MA92*/CA95^* showed surface-localized fluorescence above the background. These analyses verified site-specific labelling of Gag within the virion.

### Virus production for smFRET imaging

To produce immature HIV-1 virions with a single fluorescently labeled Gag protein, as required for smFRET imaging, we transfected each of three 10-cm plates of HEK293T cells with 19 μg pNL4-3 ΔRTΔPR, 1 μg pNL4-3 ΔRTΔPR(I92TAG/Q227TAG), and 1 μg each of Mm- PylRS-AF/Pyl-tRNACUA and eRF1 plasmids in the presence of 1mM TCO*A in the culture media. The ratio of the two pNL4-3 plasmids was empirically determined based on the observation that wild-type HIV-1 was produced *ca.* 800 times more efficiently than HIV-1^MA92*/CA95^* (see above). The media supplemented with 1 mM TCO*A was replaced 8 hours post-transfection. Virions were harvested after 48 hrs at 37°C, site-specifically labeled with JF549 and JF646, dialyzed, and purified as described above. For virions produced in the presence of 10 μM PDDC, 10 μM DPTIP, or 50 nM LEN, an identical procedure was used with the additional supplement of the inhibitors in the culture media.

### smFRET imaging with TIRF microscopy

Virions were immobilized on streptavidin-coated quartz microscope slides, as described^46^. Imaging was performed in buffer containing 20 mM Tris-HCl pH 8, 50 mM NaCl, and a cocktail of triplet-state quenchers and an enzymatic oxygen-scavenging system to stabilize fluorescence. For experiments where the virions were incubated with LEN or BVM before imaging, the imaging buffer was supplemented with the respective inhibitors. Virions were imaged on a custom-built prism based TIRF microscope. The technical details of the microscope can be found in the previous publication^37,47^. Briefly, virions were illuminated by the evanescent field generated by the total internal reflection of a 532 nm Obis solid-state laser (Coherent, Saxonburg, PA) at an intensity of 1.2 kW/cm^2^. Fluorescence emission was collected by a 1.2-NA 60X water-immersion objective (Olympus, Needham, MA). Donor and acceptor fluorescence emissions were separated by a T635LPXR dichroic filter (Chroma, Bellows Falls, VT) and imaged on parallel halves of a Prime 95B sCMOS camera (Photometrics, Tucson, AZ). 500-frame movies were acquired at 100-ms exposure using Micro-Manager 2.0.

### Analysis of smFRET data

smFRET trajectories were analyzed using SPARTAN software (https://github.com/stjude-smc/SPARTAN) with additional scripts in Matlab (Mathworks, Natick, MA). FRET was calculated using the formula *I*_A_/(*γI*_D_+*I*_A_), where *I*_A_ and *I*_D_ are the acceptor and donor fluorescence intensities, respectively, and *γ* is an empirically determined factor correcting for differences in detection efficiency. The fluorescence traces were also corrected for donor fluorescence bleadthrough into acceptor channel. smFRET trajectories were selected for analysis using the following criteria: (1) single-step photobleaching of donor and acceptor fluorescence, which indicates a single pair of donor and acceptor fluorophores per virion; (2) correlation coefficient of donor and acceptor fluorescence was less than or equal to zero; (3) minimum total fluorescence intensity was greater than mean background fluorescence intensity (210 units); (4) the average FRET value of the trace before photobleaching was greater than 0.025. smFRET trajectories that satisfied these criteria were collected and compiled into FRET histograms. The smFRET trajectories were fit to a five-state kinetic model using hidden Markov modeling (HMM) and a maximum likelihood estimation using the MPL algorithm implemented in SPARTAN. This resulted in identification of the following FRET states: 0 ± 0.025, 0.07 ± 0.03, 0.15 ± 0.05, 0.35 ± 0.09, 0.6 ± 0.12 (mean FRET efficiency ± standard deviation). Gaussian distributions reflecting these FRET states were overlaid on the FRET histograms. The results of the HMM analysis was used to determine the fraction of time each trajectory spent in each of the five FRET state. The state occupancies were compiled into violin plots in Matlab. Dwell time histograms were generated from the HMM analysis and fit to double exponential functions (*A*_1_ exp (-*t*/*τ*_1_) + *A*_2_ exp (-*t*/*τ*_2_)) to determine the average time spent in each state before transitioning. Weighted averages of the two time constants were computed and listed in Table S1. Subpopulations of smFRET trajectories was carried out using *k*-means clustering in Matlab. Clustering was performed on the following parameters: (1) combined occupancy in the 0.07- and 0.15-FRET states; and (2) the number of FRET transitions.

### MD simulations

An atomic model of an 18-mer assembly of MA-CA was generated by integrating the available coordinates of immature MA (PDB 1HIW^39^; residues 7–121) and CA (PDB 5L93^6^; residues 148–371) with an AlphaFold3-predicted model (residues 101-160) of the linker spanning the two domains. The overlapping regions of the experimental and AlphaFold3-predicted structures were aligned to produce a continuous and structurally consistent MA-CA model. Atomic models of the dyes were constructed in PyMol. Model geometries were optimized initially at the AM1 level of theory with the *sqm* program in the AmberTools software package. Geometries were further optimized and the electrostatic potential calculations were performed at the HF/6-31G(d) level of theory in Gaussian 9^37^. Partial charges were calculated by fitting restrained electrostatic potential (RESP) using the *antechamber* module from AmberTools. Atom types and bonded parameters from the Generalized Amber Force Field (GAFF2)^48^ were assigned using the *parmchk2* module from AmberTools. Dyes were attached at residues I92 of MA and Q227 of CA with LEaP. The protein was parameterized with the ff19SB Amber forcefield^49^. The system was charge-neutralized and solvated in explicit water with the OPC model^50^ with periodic boundary conditions. Additional Na+ and Cl- ions were added to achieve a salt concentration of 150 mM. Simulations of the MA-CA 18-mer and monomer were run on the SCI cluster at UMass Chan Medical School with Amber v22 using Particle Mesh Ewald Molecular Dynamics with CUDA executable for NVIDIA GPUs (PMEMD.cuda)^51^. Water and ions were energy minimized for 1000 steps. The system was heated to 298 K at a constant volume, followed by equilibration at a constant pressure with harmonic restraints on all non-solvent atoms using an initial force constant of 100 kcal/mol·Å^2^, and subsequently reduced to 10 kcal/mol·Å^2^. A second energy minimization was performed with restraints on the protein backbone and all heavy atoms of the dyes. This was followed by a constant pressure equilibration with the same restraints, which were gradually reduced from 10 to 1 and finally to 0.1 kcal/mol·Å², followed by a final equilibration with all restraints removed. MD production runs were done at a constant pressure of 1 bar with a timestep of 2 fs for 310 ns, and the first 10 ns were discarded as equilibration. Each production simulation was run in triplicate, yielding a total simulation time of 0.9 μs each for the 18-mer and the monomer. Trajectories were processed using *cpptraj* from AmberTools22^52^, and visualized in VMD. Distances between the central carbon atoms of the polymethine chain in each fluorophore was determined.

## Acknowledgements

We thank members of the Munro, Schiffer, and Freed laboratories for helpful discussions.

## Author contributions

Conceptualization: T.N., J.B.M. Methodology: T.N., J.B.M. Data acquisition: T.N., M.W. Data curation: T.N., J.B.M. Formal analysis: T.N., M.W., J.B.M. Writing – Original draft: T.N., J.B.M. Writing – Review and editing: all authors. Supervision: J.B.M., C.A.S., E.O.F.

