## Supplementary Material for "Inhibitors of HIV-1 maturation modulate conformational dynamics and heterogeneity of the immature Gag lattice"

**Supplementary figures**

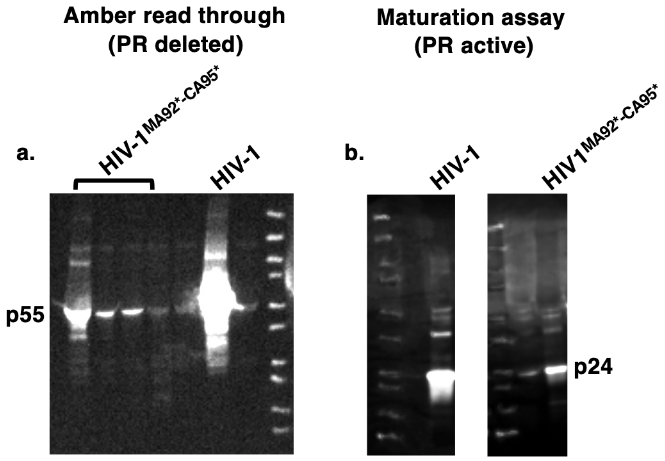


**Fig. S1. Amber read through and proteolytic maturation assay.** (**a**) p24 western blot of viral supernatant showing expression of full-length HIV-1$^{MA92*-CA95*}$ Gag and HIV-1 Gag (55 kDa), both with PR deleted. (**b, c**) p24 western blot of HIV-1 and HIV-1$^{MA92*-CA95*}$ viral supernatant showing expression of p24, resulting from proteolysis of Gag by PR. This verifies that HIV-1$^{MA92*-CA95*}$ Gag undergoes proteolytic maturation comparably to WT.

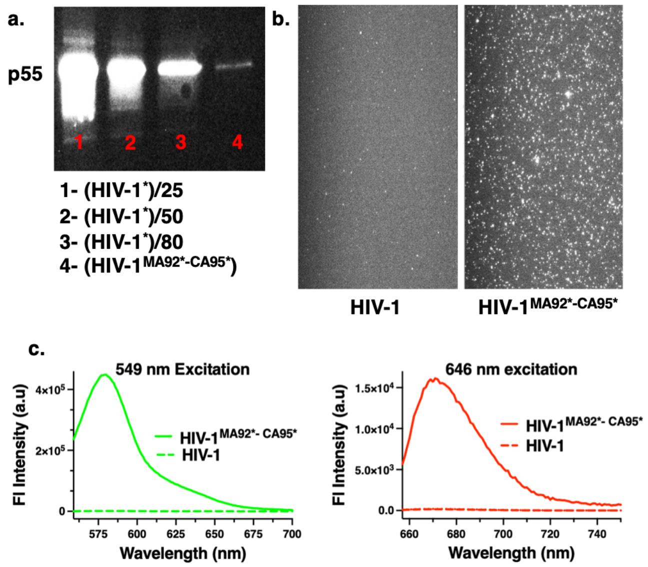


**Fig. S2. Site-specific labelling of Gag within immature HIV-1 virions.** (**a**) p24 western blot of dye labelled virions showing three dilutions of full-length HIV-1 Gag (55 kDa) and HIV-1$^{MA92*-CA95*}$. (**b**) TIRF image acquired with 532-nm excitation of (left) mock labelled HIV-1 Gag and (right) HIV-1$^{MA92*-CA95*}$ Gag. The relative concentrations of the two virus samples was normalized using the blot in (a), so that equal concentrations were surface immobilized. (**c**) Steady state fluorescence emission spectra of concentration normalized mock-labelled HIV-1 and labelled HIV-1$^{MA92*-CA95*}$. Taken together, (b) and (c) show successful site-specific labelling of HIV-1$^{MA92*-CA95*}$ and negligible labelling of wild-type HIV-1.

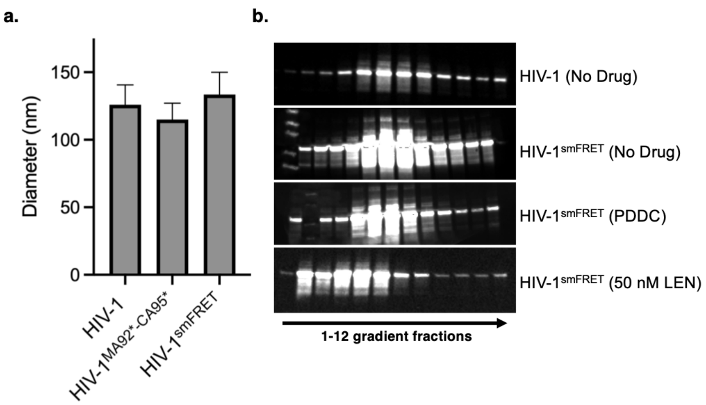


**Fig. S3. Virions formed with modified HIV-1 Gag retain native morphology.** (**a**) Size distribution of virions determined by negative stain electron microscopy shows no significant difference between HIV-1, HIV-1$^{MA92*-CA95*}$, and the smFRET sample prepared with an excess of wild-type HIV-1 over HIV-1$^{MA92*-CA95*}$ (HIV-1$^{smFRET}$). (**b**) p24 Western blot of optiprep gradient fractions showing p55, which indicates that HIV-1 and HIV-1$^{smFRET}$ migrate to the same fractions (peak fractions 4-9) after ultracentrifugation, in the absence or presence of PDDC. Virus production in the presence of 50 nM LEN leads to larger particles that migrate to fractions 2-6.

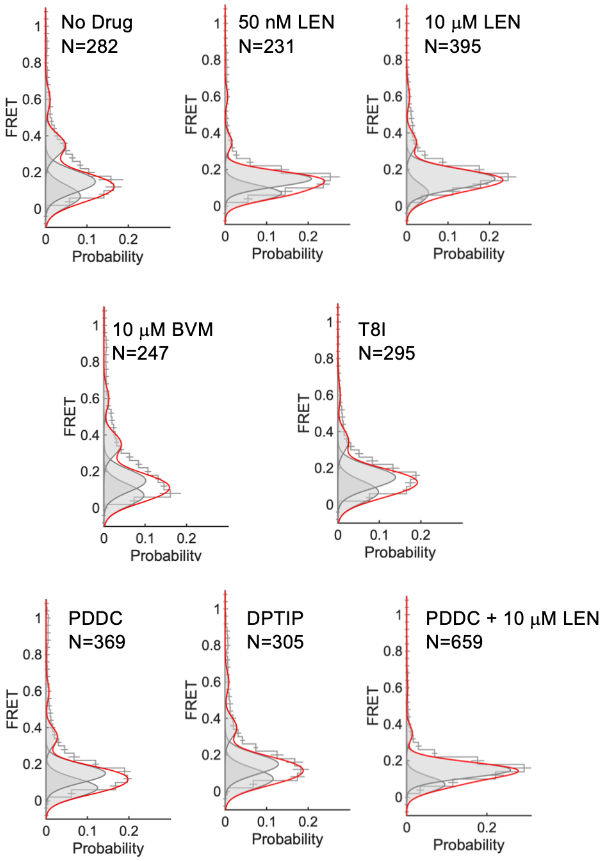


**Fig. S4. FRET histograms show Gag conformational distribution in the presence of maturation inhibitors.** smFRET data were acquired under the indicated conditions. Histograms are displayed as in Fig. 1.

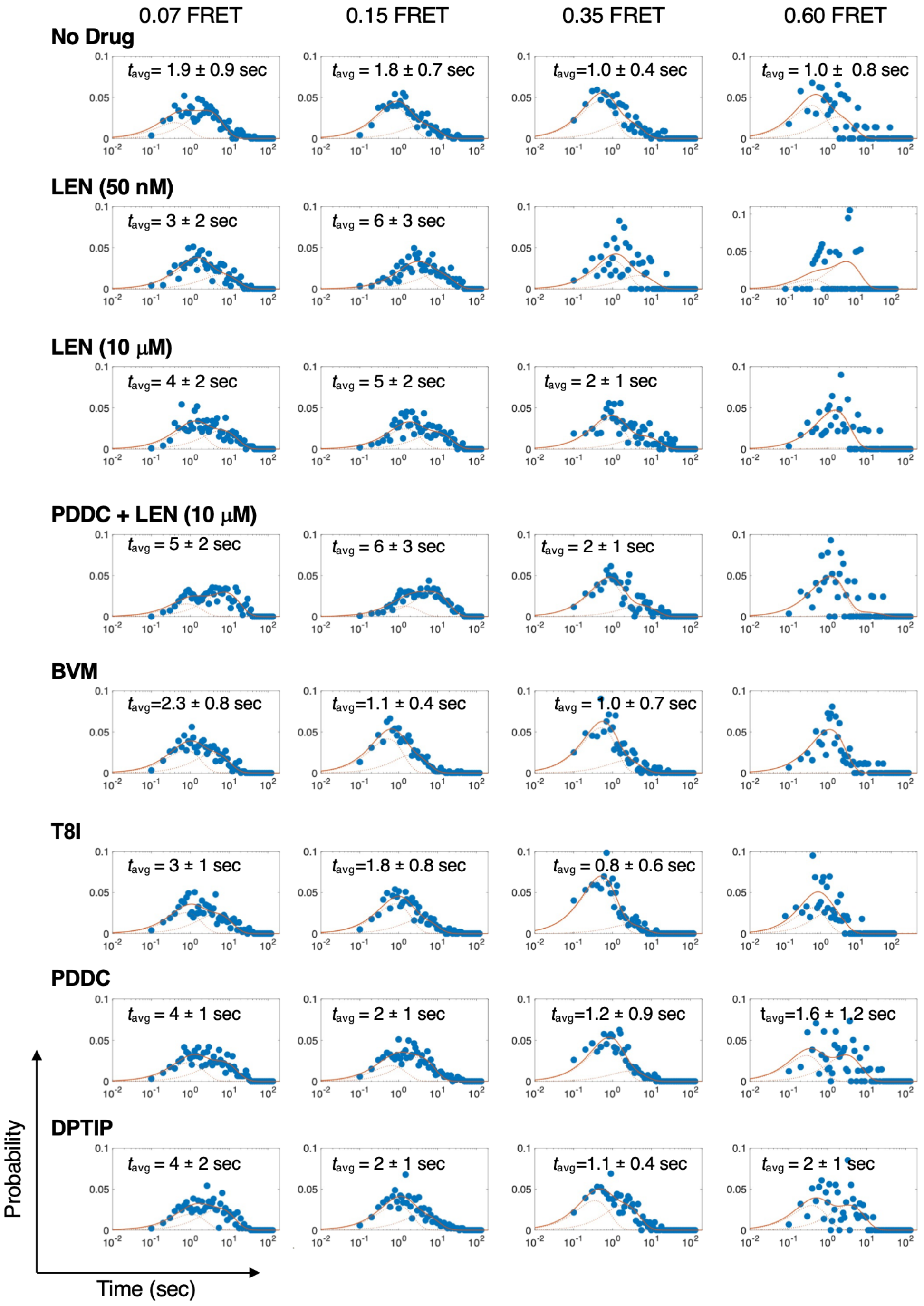


**Fig. S5. Average lifetime in each FRET state prior to transition.** Dwell time histograms of the four non-zero FRET states determined from HMM analysis. smFRET data were acquired in the presence of the indicated maturation inhibitors. All distributions are fit to a double exponential function, $A_1 e^{-t/t1} + A_2 e^{-t/t2}$, where $A_1$ and $A_2$ are the amplitudes of the two time constants, $t_1$ and $t_2$. The fits are overlaid in red and the weighted average lifetime of each state ($t_{avg} = A_1 t_1 + A_2 t_2$) is indicated ± the 95% confidence interval from the fit propagated through the weighted average calculation. Distributions are presented with logarithmically spaced bins to aid in visualization of the two exponential distributions. Where $t_{avg}$ is not indicated, it was not determined due to insufficient observations of that transition.

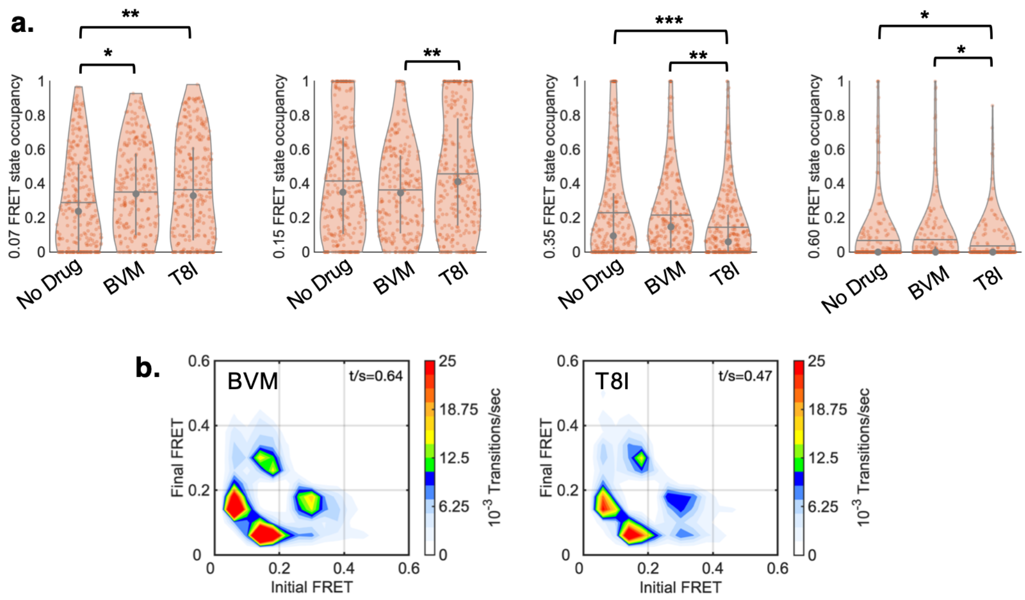


**Fig. S6. BVM and T8I stabilize the 0.07-FRET state.** (**a**) Occupancy violin plots for the four non-zero FRET states acquired in absence of inhibitor, and in the presence of BVM or the T8I mutation. Both BVM and T8I increase the 0.07-FRET state occupancy. The effect of T8I is modestly stronger than BVM. T8I further reduces 0.35 FRET state occupancy. Statistical significance is indicated with asterisks, as in Fig. 2. (**b**) TDPs acquired in the presence BVM and T8I, with transitions per second (t/s) indicated.

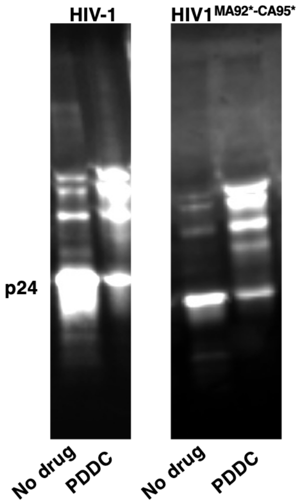


**Fig. S7. Producing virus in the presence of PDDC disrupts proteolytic maturation of HIV-1.** p24 Western blot of HIV-1 and HIV-1$^{MA92*-CA95*}$ viral supernatant (with active PR) produced in the absence or presence of PDDC. This confirms that the HIV-1$^{MA92*-CA95*}$ and HIV-1 are affected by PDDC similarly.

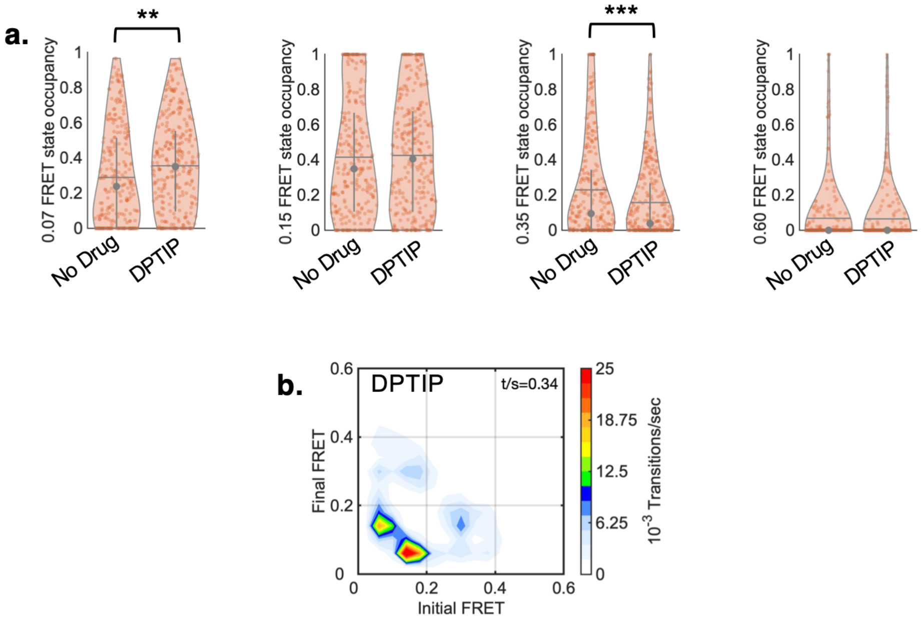


**Fig. S8. Producing virus in the presence of DPTIP stabilizes the 0.07-FRET state.** (**a**) Occupancy violin plots for the four non-zero FRET states acquired from virions produced in absence or presence of DPTIP. DPTIP increased occupancy in the 0.07-FRET state, and reduced occupancy in the 0.35-FRET state. (**b**) TDP showing reduced Gag conformational dynamics in presence of DPTIP, with transitions per second (t/s) indicated.

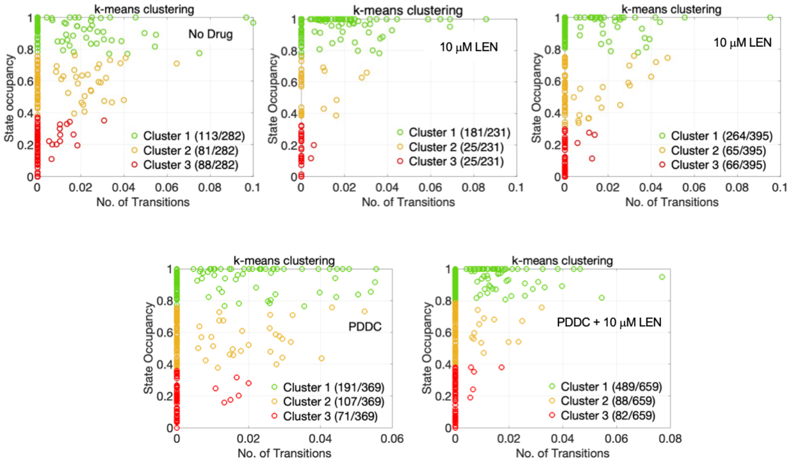


**Fig. S9. Clustering of FRET trajectories is consistent with Gag lattice heterogeneity within the immature virion.** *k*-means clustering of smFRET traces was based on the combined state occupancies of the 0.07- and 0.15-FRET states, and number of transitions. The number of trajectories in each cluster is indicated in the numerator, with the denominator indicating the total number of trajectories. The Cluster percentages are shown in Fig. 4.

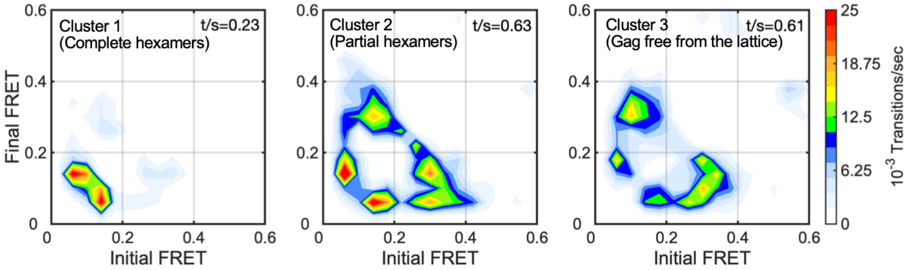


**Fig. S10. FRET trajectories within each Cluster display distinct conformational dynamics.** TDPs of the three individual clusters determined in the absence of maturation inhibitors. t/s indicates the number of transitions per second.
